# Ethylene biosynthesis in guard cells, not mesophyll, predominantly drives stomatal conductance responses to CO_2_

**DOI:** 10.64898/2026.03.05.708972

**Authors:** Dikla Nissan-Roda, Or Shapira, Danielle Neta, Shira Gal, Tamar Azoulay-Shemer

## Abstract

- **Research and rationale**: This study investigates whether tissue-specific ethylene biosynthesis regulates stomatal conductance (g_s_) responses to changing [CO₂] in *Arabidopsis thaliana*. While guard cells sense [CO₂], mesophyll-derived signals are also implicated in stomatal control. We aimed to determine if ethylene production in guardcells or mesophyll is the primary driver of CO₂-induced g_s_ regulation.
- Methods: An *acs* octuple mutant with severely reduced ethylene production was complemented with tissue-specific *ACS8/ACS11* transgenes driven by guard-cell, spongy-mesophyll, dual palisade/spongy-mesophyll, or whole-leaf promoters. Tissue-specific complementation in the different transgenic lines was confirmed and evaluated by qPCR, tissue-specific NEON expression, microscopic imaging, and ethylene production measurements. Gas-exchange measurements on intact plants recorded g_s_ kinetics, CO_2_ assimilation, and water-use efficiency, across CO₂ shifts.
- **Key results**: Guard-cell complementation nearly fully restored wild-type g_s_ responses and reversed the mutant’s aberrant leaf phenotype. Spongy-mesophyll complementation failed to rescue either trait, while dual palisade- and spongy-mesophyll complementation yielded only partial recovery.
- **Conclusion:** Ethylene produced in guard cells is the dominant regulator of CO₂-induced stomatal conductance regulation, with mesophyll-derived ethylene contributing secondarily via long-distance signaling or by augmenting the overall ethylene pool. These findings underscore the importance of spatially regulated ethylene biosynthesis in balancing carbon assimilation and transpiration.

## Introduction

Land plants mediate gas exchange between the atmosphere and the interior of the leaf by tightly regulating stomatal conductance to assimilate CO_2_ for photosynthesis and simultaneously prevent excessive water loss. The complex regulatory mechanisms of the stomata are crucial for the optimal response of the plant to the environment, for plant physiology, and best plant performance. Stomata, found on the surface of above-ground plant tissues, are tiny pores constructed by a pair of guard cells that increase and decrease in size to control the gap between them, thereby controlling the rate of transpiration and the diffusion of CO_2_ (Hetherington & Woodward, 2003; Yi *et al*., 2022). Different environmental factors, tightly regulate the opening and closing of the stomata, including drought (Schroeder *et al*., 2001b; Daszkowska-Golec & Szarejko, 2013), ozone (Hasan *et al*., 2021), humidity (Merilo *et al*., 2018; Jalakas *et al*., 2021), light (Taylor *et al*., 2024; Zhu *et al*., 2025), pathogens (Maidment & Xu, 2025), and CO_2_ (Engineer *et al*., 2016; Dubeaux *et al*., 2021), all acting through interconnected signaling pathways in guard cells and adjacent tissues (Bawa *et al*., 2023), integrative regulatory mechanisms of stomatal movements in response to environmental stresses (Zhang *et al*., 2024).

The mechanisms of the guard cell for sensing stimuli, including CO_2_, are well-established (Tian *et al*., 2015; Engineer *et al*., 2016; Dubeaux *et al*., 2021; Takahashi *et al*., 2025). Several studies have provided evidence supporting the presence of mesophyll-driven signals that move from the mesophyll to the epidermal layer, contributing to the regulation of stomatal conductance (Wong *et al*., 1979; Lee & Bowling, 1992; Lake *et al*., 2002; Mott, 2009; Fujita *et al*., 2013; Fujita *et al*., 2019; Santos *et al*., 2021). It has been suggested that these signals include both soluble (Fujita *et al*., 2013) and vapor-phase signals (Sibbernsen & Mott, 2010; Mott *et al*., 2014).

Ethylene, a gaseous plant hormone, is involved in various stress-induced stomatal conductance regulation responses, such as ozone, drought, and ABA (Dodd, 2003; Pospíšilová, 2003; Tanaka *et al*., 2005; Tanaka *et al*., 2006; Wilkinson & Davies, 2009).

CO_2_, one of the primary regulators of stomatal conductance, is sensed by the guard cells in the sub-stomatal cavity (Ci). Elevated [Ci] in leaves results in a reduced stomatal conductance expressed in stomatal closure, while reduced [Ci] results in stomatal opening (Vavasseur & Raghavendra, 2005; Tian *et al*., 2015; Engineer *et al*., 2016). The regulation of ethylene production by [CO_2_], which is inconsistent among different studies, suggests different regulatory modes by CO_2_ in various tissues and environmental conditions (Mathooko, 1996; Ahammed & Li, 2022). Yet, the effect of ethylene on CO₂-induced stomatal movements remains unclear. Ethylene has been reported to induce stomatal opening in some studies (Iqbal *et al*., 2011; Gautam *et al*., 2022), whereas others showed ethylene-mediated stomatal closure (Desikan *et al*., 2006). More recently, ethylene biosynthesis and signaling have been shown to modulate the kinetics and sensitivity of stomatal conductance responses to CO₂ and ABA (Azoulay-Shemer *et al*., 2023). Although few studies have questioned ethylene’s role in stomatal conductance regulation, analyses were mainly done on epidermal peels or isolated guard cells (Desikan *et al*., 2006), and very few studies used intact plants (Azoulay-Shemer *et al*., 2023). The role ethylene plays in stomatal conductance regulation as long-distance signals from the mesophyll versus autonomously within the guard cells are brought in this study.

As sessile organisms, plants have evolved different mechanisms to respond to their ever-changing environments. Communication among other parts/tissues of the plant is essential for the plant to respond appropriately to changes in its surroundings (Baluska, 2013; Huber & Bauerle, 2016; Johns *et al*., 2021). Mesophyll is the primary photosynthetic and carbon-metabolizing tissue in C3 plants. It is located between the underside (abaxial side) and the upper (adaxial) epidermal layers, including the guard cells, which regulate gas exchange between the atmosphere and the leaf’s interior. In dicots, such as Arabidopsis, xll consists of spongy mesophyll, which forms intercellular spaces for the diffusion of CO_2_ from stomatal pores on the underside (abaxial side) of the leaf (Borsuk *et al*., 2022)and compactly organized palisade cells, surrounded by apoplastic fluid, toward the upper (adaxial) side of the leaf (Trozzi & Robil, 2025).

Various mechanisms tightly regulate ethylene biosynthesis (Booker & DeLong, 2015; Pattyn *et al*., 2021) in a tissue-dependent matter (Vong et al., 2019). Ethylene biosynthesis has been shown to depend on the expression and activity of the ACC synthase enzymes, a rate-limiting step in ethylene production (Yang & Hoffman, 1984; Khan *et al*., 2024; Li *et al*., 2025). In Arabidopsis, there are nine functional ACS enzymes (out of 12 encoded genes), which are differentially regulated (Tsuchisaka & Theologis, 2004; Tsuchisaka *et al*., 2009) and are expressed in different tissues, including guard cells, vascular tissue, and mesophyll (Yamagami *et al*., 2003; Peng *et al*., 2005). The distinctive temporal and spatial expression patterns of these genes (Datta *et al*., 2015) and the post-translational regulation of these enzymes (Yoon, 2015) control ethylene production in a tissue-specific manner (Vaseva *et al*., 2016).

This study aims to identify the source tissue/cells that produce ethylene and is/are involved in CO_2_-induced stomatal movement. In a previous study by Tsuchisaka *et al*. (2009), a high-order *acs* octuple Arabidopsis mutant was generated, defective in eight out of the nine active ACS genes. A detailed phenotypic work showed severely reduced levels of ethylene (under ambient [CO_2_]) in this mutant (Tsuchisaka *et al*., 2009; Azoulay-Shemer *et al*., 2023). Gas exchange measurements revealed that the *acs* octuple mutant exhibits dysfunctional CO_2_-induced stomatal movements compared to wild-type (WT, Col-0) plants, which supports ethylene involvement in stomatal conductance responses to [CO_2_] (Azoulay-Shemer *et al*., 2023).

In this research, we analyze the role of tissue-specific ethylene biosynthesis in stomatal conductance by complementing the *acs* octuple mutant in a tissue-specific matter; in the stomata guard cells, the spongy mesophyll cells, the spongy and palisade mesophyll cells, and the whole plant. Gas exchange measurements and ethylene production quantification revealed that both mesophyll and guard cell ethylene biosynthesis contribute to stomatal conductance regulation, with guard cell-derived ethylene playing a primary role. These results highlight the importance of localized ethylene signaling in optimizing plant responses to environmental changes.

## Materials and Methods

### Plant materials and mutant lines

This study used the Arabidopsis (*Arabidopsis thaliana*) plant model. Columbia (Col-0) ecotype was used as the wild-type (WT) for comparison. The *acs* octuple mutant, a high-order mutant in eight genes of the ACC synthase family (cs16651: *acs* sextuple mutant silenced in *ACS8* and *ACS11* (amiR)) mutants (Tsuchisaka *et al*., 2009), was ordered from ABRC.

### Plant growth conditions

Soil-grown plants*: Arabidopsis thaliana* seeds were surface-sterilized and germinated on 0.5 MS medium (Murashige & Skoog, 1962) plus 0.8% (w/v) phyto agar and 0.8% (w/v) sucrose or on autoclaved soil (Even Ari Green 761 Arabidopsis Blending) + 2g/L Osmocote®. Seeds were stratified for two days at 4°C and then moved to the growth room for germination. 10 to 15-day-old seedlings were transferred to 250 ml soil containing pots, two seedlings per pot, and watered twice weekly. All Arabidopsis plants were grown in a plant control growth room or Percival growth chambers under uniform environmental conditions, under a 12-h light (7:00–19:00) /12-h dark photoperiod at 19°C/21°C (respectively), ∼150-180μmol m^−2^ s^−1^ photon flux density, and relative humidity of 60-80%.

### Gas exchange measurements

Gas exchange measurements were performed on the fifth/sixth true leaf of 5-6.5 weeks-old intact plants. Data was logged 1-3 hours after the light turned on using a Li-6800 infrared (IRGA)-based gas exchange analyzer system equipped with a fluorimeter chamber (LI-COR Biosciences, Lincoln, NE, USA). During measurement, the Li-6800 leaf chamber was set to an air temperature of 21°C, relative humidity of 60%, and a photon flux density of 250 µmolm^-2^ sec^-1^ (with 10% blue light). Throughout the experiments, Wild-type (WT, Col-0) and mutant plants were grown in parallel under the same growth conditions.

Stomatal responses to shifts in [CO_2_] were conducted as follows: stomatal conductance was first stabilized at ambient CO_2_ levels (e.g., 360/415 ppm) for 15 minutes, as indicated in each figure. Subsequently, [CO_2_] was shifted to 800/900 ppm for 20 minutes, followed by a further shift to 100 ppm for an additional 50 minutes. The figure legends provide details regarding the number of biological replicates (n = number of plants) and the number of stomata analyzed in each sample.

### Statistical analysis

Statistical analysis was performed using GraphPad Prism software version 10.0 (GraphPad Software, San Diego, CA, USA, www.graphpad.com). Data was checked for normality of the residuals around the mean and for equal variance using the Shapiro-Wilk and Brown-Forsyte tests, respectively. Means between treatments were compared using Student’s t-test for single comparison, or Tukey’s multiple comparisons test following One-way ANOVA. Stomatal closing and opening rates were calculated as the slope of stomatal conductance kinetics in response to changes from ambient to high (first 5 minutes, closing) and high to low CO_2_ concentration (first 20 minutes, opening). The slope was determined using linear regression and compared using their 95% confidence intervals. Asterisks indicate significant differences between genotypes as *P* < 0.03 (*); 0.002 (**); 0.001 (***).

### Complementation of the *acs* octuple line

To achieve tissue-specific complementation in the *acs* octuple mutant, we cloned the modified versions of *ACS8* and *ACS11* genes under the following specific promoters:

1. <u>The guard cell-specific promoter, *GC1* (AT1G22690)</u> (e.g., GCD1) (Yang *et al*., 2008) sequence, was kindly provided by Prof. Julian Schroeder (UCSD, California).
2. <u>The spongy mesophyll-specific promoter *CORONATINE INDUCED 3* (AT4G23600)</u> (e.g., CORI3) (Procko *et al*., 2022).
3. <u>The palisade mesophyll *IQ-DOMAIN 22 (IQD22)* (AT4G23060) promoter (e.g., PEG1)</u>- (Geisler *et al*., 2002). Both CORI3 and PEG1-specific promoters were kindly provided by Dr. Carl Procko and Joanne Chory (Salk Ins, CA, USA).
4. <u>*UBIQUITIN 10* (AT4G05320)</u> is the whole-plant expression promoter (e.g., UBQ10) (Sun & Callis, 1997).

For schematics of DNA constructs and expression promoters, see Methods S1.

All constructs were generated using the “Golden Gate” multi-cassette cloning strategy (Engler *et al*., 2014). Plasmid construction and primer design were performed in the online *Benchling* software (https://www.benchling.com/). The primer sequences used for constructing the L0 level are detailed in Table S1. For fast screening and detection of positive transformants transgenic lines, each construct includes another cassette expressing the NEON fluorescent protein under the same tissue-specific promoter and the VENUS fluorescent protein (Nagai *et al*., 2002) under the seed coat-specific expression promoter. The target vectors were introduced into the *acs* octuple mutant genome by the *Agrobacterium*-mediated transformation method (Bechtold, 1995). Seeds were collected from the *acs* octuple transformant (T0) and screened for positive transformants under a fluorescence stereo microscope (Leica M205 FA) based on VENUS seed coat protein fluorescence. The fluorescent seeds were germinated, and their leaves were screened using a light microscope (Olympus BX61 Upright Microscope) for tissue-specific NEON expression. Next, the different transgenic lines were verified by PCR using specific primers (see Table S2). At least three positive lines for each construct were isolated for self-fertilization and seed propagation. Confocal microscopy was conducted on the T1 generation plants at the Volcani Center microscopy unit for further investigation. T2 generation plants were used for CO_2_ gas exchange analysis.

### Quantitative RT-PCR analysis

Total RNA was extracted using the plant total RNA purification kit (Norgen Biotek Corp, Canada) following the manufacturer’s instructions. First-strand cDNA was synthesized from ⁓400 ng of total RNA for real-time PCR analysis using UltraScript™ cDNA Synthesis Kit (PCR Biosystems, UK). Real-time PCR was performed using the Applied Biosystems™ StepOne™ Real-Time PCR System (Applied Biosystems) with SYBR™ Green PCR Master Mix (Applied Biosystems) in the amplification mixture according to the manufacturer’s protocols. Primer sets for each gene are listed in Table S3. The amplification of the actin transcript served as the internal standard, and the data were analyzed with StepOnePlus™ Software v2.3 (Applied Biosystems). The relative transcript levels of *ACS8* and *ACS11* genes were calculated using the 2^-ΔΔCT^ method (Livak & Schmittgen, 2001) and standardized with the *Atactin8* transcript level.

### Microscope analysis

All fluorescence and stomatal imaging experiments were performed on a light microscope, Olympus BX61 Upright Microscope, with a DP73 digital camera. For stomatal development analysis, imaging was performed using the differential interference contrast (DIC) at 40× magnification (0.20 mm^-2^ field of view). The fluorescence stereo microscope (Leica M205 FA) and Leica DFC360 FX monochrome digital camera were used to screen for fluorescent seed signals. Confocal microscopy of fluorescent-labeled cells was done using a Leica SP8 laser scanning microscope (Leica, Wetzlar, Germany), equipped with a solid-state laser with 488nm light, HC PL APO 10×/0.4 objective, HC PL APO CS2 20×/0.75 objective or HC PL APO CS 63×/1.2 water immersion objective (Leica, Wetzlar, Germany) and Leica Application Suite X software (LASX, Leica, Wetzlar, Germany). Imaging of the GFP signal was done using the 488 nm laser line excitation, and the emission was detected in a range of 495–545 nm.

### Quantification of ethylene production

Ethylene production was quantified in 4–to 5-week-old plants grown under ambient CO_2_ conditions. Aerial plant parts were excised and placed in 15 ml Falcon tubes promptly plugged with a rubber cap. Ninety minutes later, ethylene was measured by gas chromatography (5890A Hewlett Packard Gas Chromatograph; Agilent Technologies, Santa Clara, CA, USA) using a short 1-ml column (13018-U, 80/100 μm Hayesep Q; Supelco; Sigma-Aldrich Inc) with flame ionization detection and calculated curve reading using a standard (Schmelz *et al*., 2003; Azoulay-Shemer *et al*., 2023).

## Results

### Complementation of *ACS8* and *ACS11*, specifically in spongy mesophyll cells, does not reverse the *acs* octuple mutant’s defective plant phenotype, nor its impaired stomatal conductance responses to [CO_2_]

Transgenic Arabidopsis plants, *CORI3::ACS8/11,* which complement the *acs* octuple mutant by expressing the *ACS8* and *ACS11* genes under the spongy mesophyll (*CORI3*) promoter (Procko *et al*., 2022), were generated. Following PCR-based screening (using the binary specific vector’s primers), three independent positive *CORI3::ACS8/11* lines (e.g., #14-14, #16-1, and #31-1) were selected for detailed physiological and molecular investigations.

#### Plant phenotype

Five-week-old Arabidopsis plants of WT, *acs* octuple, and three different *CORI3::ACS8/11* transgenic lines were examined for their anatomical phenotype. Results indicate an aberrant leaf phenotype in the *acs* octuple mutant, with a thinner leaf blade and a downward curling tip compared to WT, as previously reported by (Tsuchisaka *et al*., 2009). In all three *CORI3::ACS8/11* transgenic lines, the *acs* octuple mutant phenotype has been conserved (Fig. 1 A-1 & S1 A-1). Only *CORI3::ACS8/11* line #16-1 exhibited slightly smaller leaves when compared to the *acs* octuple mutant plant (Fig. S1 A-1).

**Figure 1:**
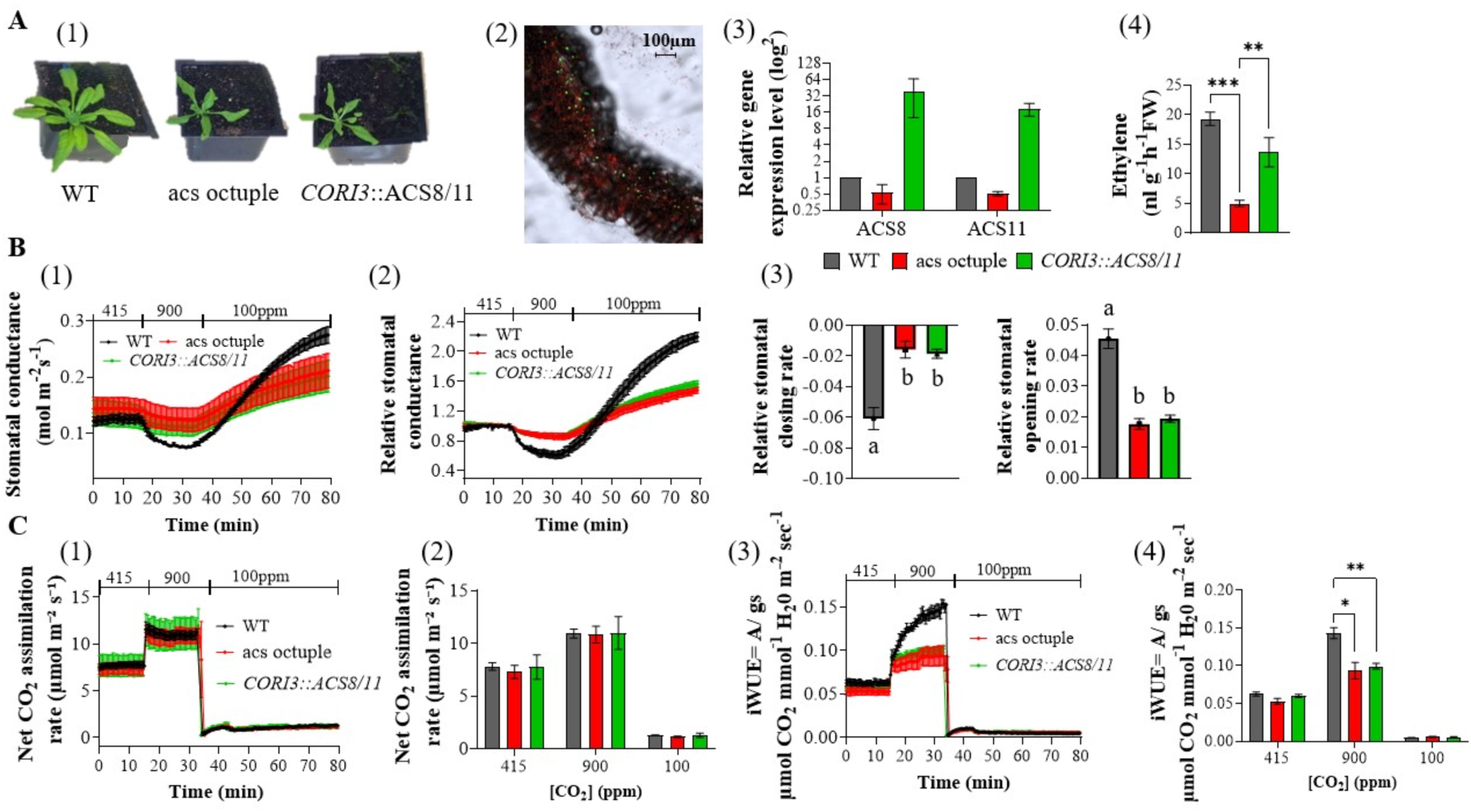
Physiological and molecular characterization of *CORI3::ACS8/11* transgenic line (#14-14): 5-week-old WT (Col-0), the *acs* octuple mutant, and the *CORI3::ACS8/11* transgenic Arabidopsis plants (line *#14-14),* which express the *ACS8 and ACS11* under the spongy mesophyll cell-specific promoter CORI3, were analyzed for the following measurements: **(A)** 1) Plant phenotype 2) Confocal microscopy analysis of the NEON green fluorescent protein, which is expressed in the spongy mesophyll cells under the CORI3 promoter. The size bar on the bottom left represents 100µm. 3) qRT-PCR leaf gene expression levels of *ACS8* and *ACS11*. Data were normalized to the WT *ACT8* gene expression level. Results are presented as relative gene expression levels (log^2^) ±SEM, n=3 plants per genotype; *P*<0.05, two-way ANOVA followed by Fisher’s LSD test. 4) Ethylene production levels *of* plant rosette leaves ±SEM of WT (n=6), *acs* octuple (n=3) and the *CORI3::ACS8/11* transgenic line *#14-14* (n=3); *P*<0.05, one-way ANOVA followed by Dunnett test. **(B)** Time-resolved stomatal conductance responses were analyzed at the imposed [CO_2_] -shifts (indicated at the top in ppm) in intact plant leaves of WT, *acs* octuple, and *CORI3::ACS8/11*. 1) Stomatal conductance in mol H_2_O m^−2^ sec^−1^. 2) Relative stomatal conductance. Data were normalized to the stomatal conductance at the last 15-sec time-point under 415 ppm [CO_2_] before [CO_2_] was shifted to 900 ppm. 3) Relative stomata closing and opening rates were calculated with a linear regression line (±SEM). The line was calculated between the 5^th^ and 20^th^ minute, under high or low [CO_2_], respectively, and the linear least squares regression statistical fitting method was used. A 95% confidence interval was used for means comparison within each line. Groups with different letters are significantly different (*P*<0.05) based on non overlapping 95% confidence intervals. **(C)** 1) Net CO_2_-assimilation rates (μmol CO_2_ m^−2^ sec^−1^). (2) The averaged net assimilation rate was calculated as the mean of the final data point for each CO_2_ level (i.e., 415, 900, 100ppm). (3) Intrinsic water use efficiency (iWUE). (4) Averaged intrinsic water use efficiency was calculated as the mean of the final data point for each CO_2_ level. Data are the mean (±SEM) of n=5 leaves from individual plants per genotype. Asterisks indicate significant differences between genotypes as *P* < 0.03 (*); 0.002 (**); 0.001 (***). Comparable findings were made in additional independent experiments, using two independent *CORI3::ACS8/11* transgenic lines (Supporting Information Fig. S1).

#### Confocal microscopy

The CORI3 promoter regulates gene expression, specifically in the spongy mesophyll (Procko *et al*., 2022). To evaluate the tissue-specificity and the level of expression mediated under the CORI3 promoter, we included in the *CORI3::ACS8/11* plasmid an additional cassette that encodes the NEON green fluorescent reporter protein under the CORI3 promoter (provided in Methods S1). Confocal analysis of the leaf cross-section of all three *CORI3*::*ACS8/11* transgenic lines showed that they express the NEON reporter protein within the spongy mesophyll cells (Fig. 1 A-2).

**Figure 2:**
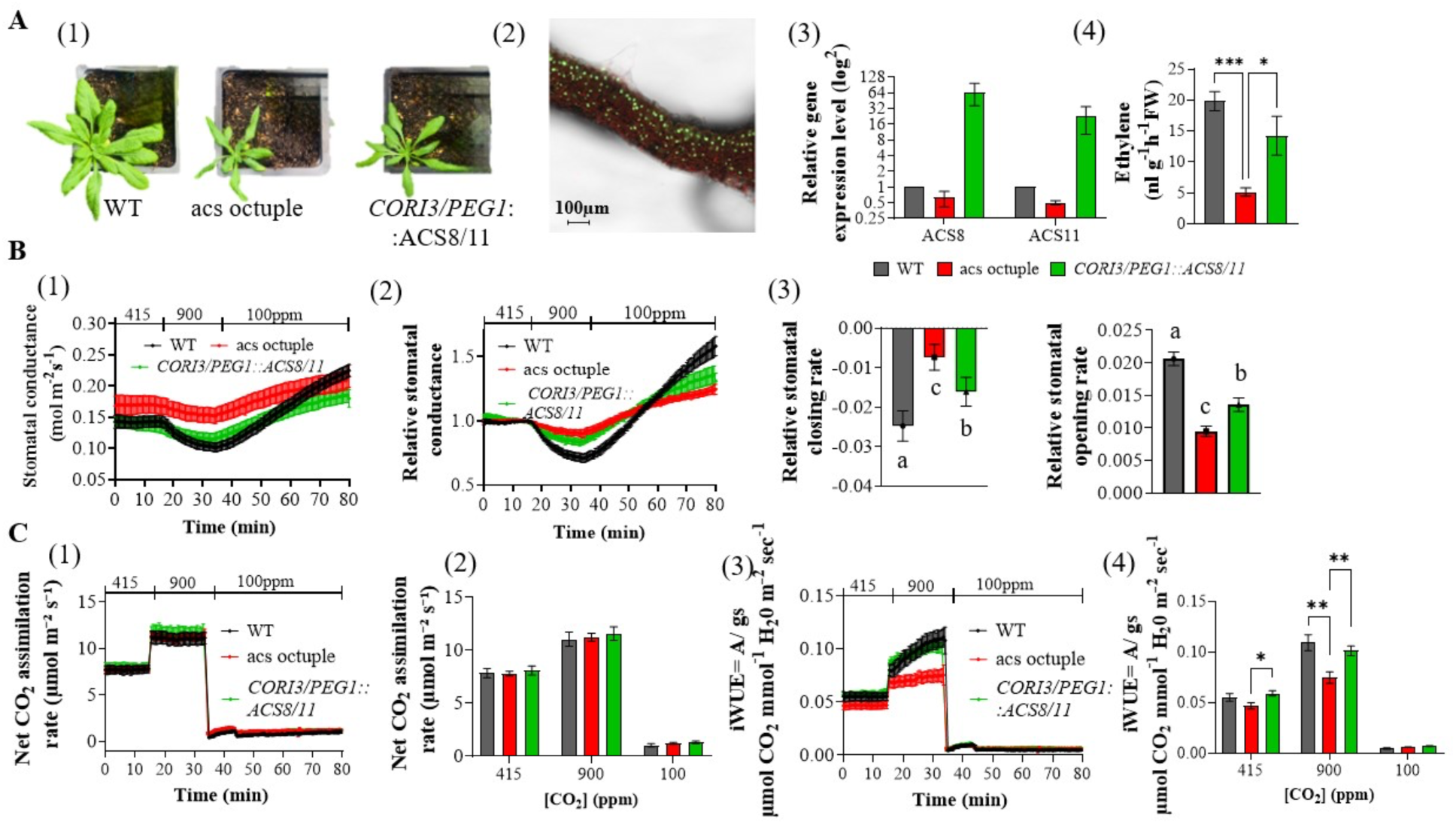
Physiological and molecular characterization of *CORI3/PEG1::ACS8/11* transgenic line (#6-2): 5-week-old WT, *acs* octuple mutant, and the *CORI3/PEG1::ACS8/11* transgenic Arabidopsis plant (line *#6-2)* expressing the *ACS8 and ACS11* under the spongy mesophyll cell-specific promoter, *CORI3,* and the palisade mesophyll, *PEG1*. The line was analyzed for the following measurements: **(A)** 1) Plant phenotype, 2) Confocal microscopy analysis of the NEON green fluorescent protein, expressed in the spongy and palisade mesophyll cells under the *CORI3 and PEG1* promoter. The size bar on the bottom left represents 100µm. 3) qRT-PCR leaf gene expression levels of *ACS8* and *ACS11*. Data were normalized to the WT *ACT8* gene expression level. Results are presented as relative gene expression levels (log^2^) ±SEM, n=3 plants per genotype; *P*<0.05, two-way ANOVA followed by Fisher’s LSD test. 4) Ethylene production levels of *plant rosette* leaves ±SEM of WT (n=6), *acs* octuple (n=6) and the *CORI3/PEG1::ACS8/11* transgenic line *#6-2 (n=3)*; *P*<0.05, one-way ANOVA followed by Dunnett test. **(B)** Time-resolved stomatal conductance responses were analyzed at the imposed [CO_2_]-shifts (indicated at the top in ppm) in intact plant leaves of WT, *acs* octuple, and *CORI3/PEG1::ACS8/11*. 1) Stomatal conductance in mol H_2_O m^−2^ sec^−1^. 2) Relative stomatal conductance. Data were normalized to the stomatal conductance at the last 15-sec time-point under 415 ppm [CO_2_] before [CO_2_] was shifted to 900 ppm. 3) Relative stomatal closing and opening rates were calculated with a linear regression line (±SEM). The line was calculated between the 5^th^ and 20^th^ minute, under high or low [CO_2_], respectively, and the linear least squares regression statistical fitting method was used. A 95% confidence interval was used for means comparison within each line. Groups with different letters are significantly different (*P*<0.05) based on non overlapping 95% confidence intervals. **(C)** 1) Net CO_2_-assimilation rates (μmol CO_2_ m^−2^ sec^−1^). (2) The averaged net assimilation rate was calculated as the mean of the final data point for each CO_2_ level (i.e., 415, 900, 100ppm). (3) Intrinsic water use efficiency (iWUE). (4) Averaged intrinsic water use efficiency was calculated as the mean of the final data point for each CO_2_ level. Data are the mean (±SEM) of n=5 leaves from individual plants per genotype. Asterisks indicate significant differences between genotypes as *P* < 0.03 (*); 0.002 (**); 0.001 (***). Comparable findings were made in additional independent experiments, using two independent *CORI3/PEG1::ACS8/11* transgenic lines (Supporting Information Fig. S2).

#### *ACS8* and *ACS11* expression in the leaf by qRT-PCR

To further validate the complementation of the *acs* octuple mutant within the spongy mesophyll cells, we extracted RNA from leaves of 5-weeks-old WT, *acs* octuple, and the three *CORI3::ACS8/11* transgenic lines. qRT-PCR analysis showed low expression levels of *ACS8* and *ACS11* genes in the *acs* octuple mutant compared to their expression levels in WT (Col-0) plants. Fig. 1 A-3 shows the relative gene expression level mean difference ± SEM and *P-*value of the effect size (*P*_ES_): ***ACS8*** -0.5± 0.2; *P*_ES WT – *acs*_ = 0.16; ***ACS11*** -0.5± 0.06; *P*_ES WT – *acs*_ = 0.08. qRT-PCR analyses of the three transgenic lines (#14-14, #16-1, and #31-1) revealed significantly high expression levels of both *ACS8* and *ACS11* compared to their expression levels in the *acs* octuple mutant. Fig. 1 A-3 shows the relative gene expression level mean difference ±SEM and *P-*value of the effect size (*P*_ES_) of line #14-14: ***ACS8*** -38.9± 26.8; *P*_ES #14-14 – *acs*_ = 0.28; *ACS11* - 17.9± 4.5, *P*_ES #14-14 – *acs*_ = 0.15. Supplemental table S4 presents raw data for the different *CORI3::ACS8/11* transgenic lines.

**Figure 3:**
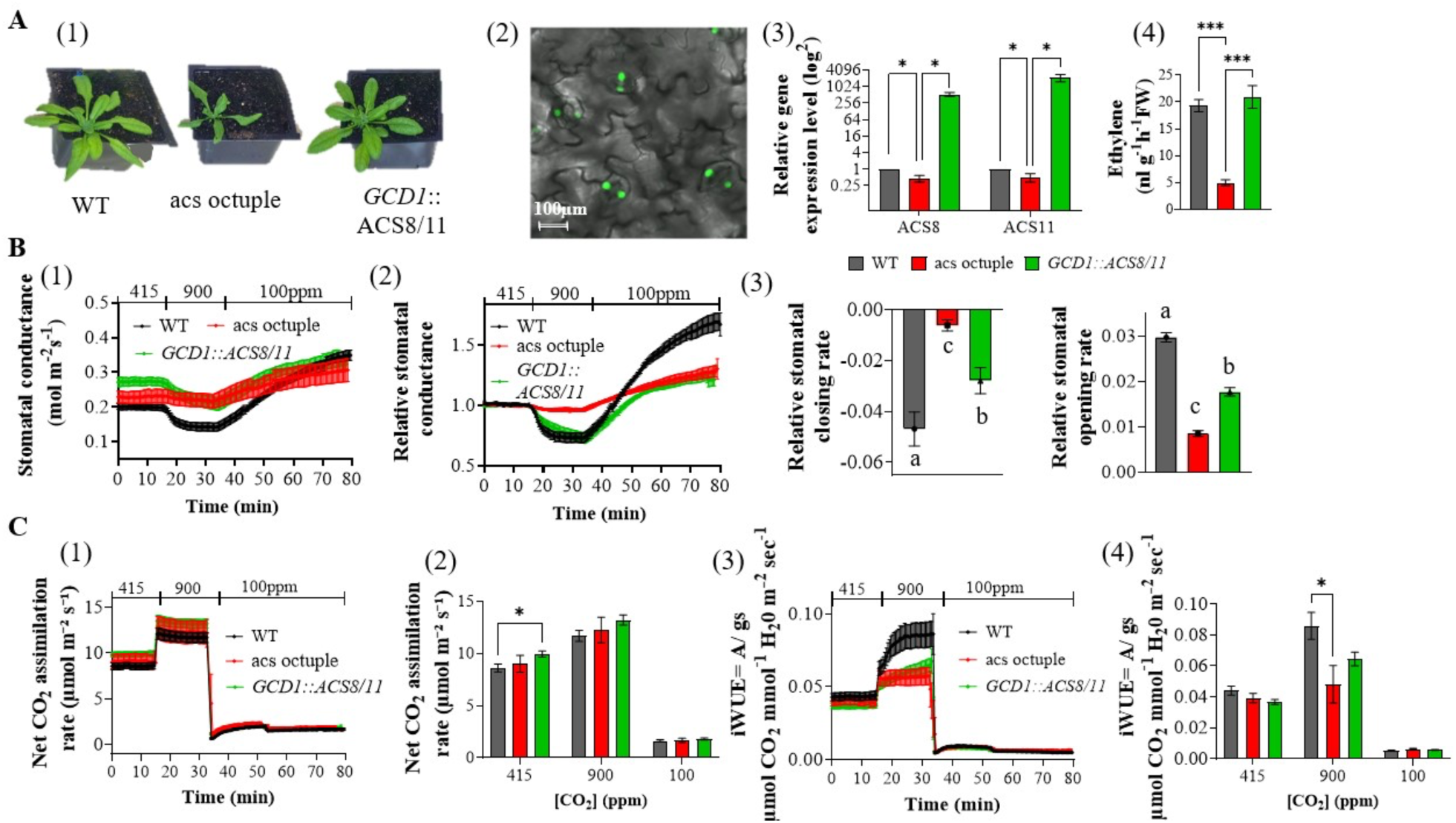
Physiological and molecular characterization of *GCD1::ACS8/11* transgenic line: 5-week-old WT (Col-0), the *acs* octuple mutant, and the *GCD1::ACS8/11* transgenic Arabidopsis plant (line *#8-5)* express the *ACS8 and ACS11* under the guard cell-specific promoter GCD1 were analyzed for the following measurements: **(A)** 1) Plant phenotype 2) Confocal microscopy analysis of the NEON green fluorescent protein, which is expressed under the GCD1 promoter. The size bar on the bottom left indicates 20µm. 3) qRT-PCR of *ACS8* and *ACS11* genes expressed in guard cells from enriched epidermal peel leaf samples. Data were normalized to the WT *ACT8* gene expression level. Results are presented as relative gene expression levels (log^2^) ±SEM (n=3) plants per genotype; *P*<0.05, two-way ANOVA followed by Fisher’s LSD test. 4) Ethylene production levels of whole plant rosettes ±SEM (n=7/3 plants for WT and *acs* octuple/ line *#8-5*; *P*<0.05, one-way ANOVA followed by Dunnett test). **(B)** Time-resolved stomatal conductance responses were analyzed at the imposed [CO_2_]-shifts (indicated at the top in ppm) in intact plant leaves of WT, *acs* octuple, and *GCD1::ACS8/11*) Stomatal conductance in mol H_2_O m^−2^ sec^−1^ 2) Relative stomatal conductance. Data were normalized to the stomatal conductance at the last 15-sec time-point under 415 ppm [CO_2_] before [CO_2_] was shifted to 900 ppm. 3) Relative stomata closing and opening rates were calculated with a linear regression line (±SEM). The line was calculated between the 5^th^ and 20^th^ minute, under high or low [CO_2_], respectively, and the linear least squares regression statistical fitting method was used. A 95% confidence interval was used for means comparison within each line. Groups with different letters are significantly different (*P*<0.05) based on non overlapping 95% confidence intervals. **(C)** 1) Net CO_2_-assimilation rates (μmol CO_2_ m^−2^ sec^−1^). (2) The averaged net assimilation rate was calculated as the mean of the final data point for each CO_2_ level (i.e., 415, 900, 100ppm). (3) Intrinsic water use efficiency (iWUE). (4) Averaged intrinsic water use efficiency was calculated as the mean of the final data point for each CO_2_ level. Data are the mean (±SEM) of n=5 leaves from individual plants per genotype. Asterisks indicate significant differences between genotypes as *P* < 0.03 (*); 0.002 (**); 0.001 (***). Comparable findings were made in additional independent experiments using two independent *GCD1::ACS8/11* transgenic lines (Supporting Information Fig. S3).

#### Ethylene production

In an earlier study, ethylene production of the *acs* octuple mutant plant leaves was found to be impaired, reaching ∼26% of WT levels (Azoulay-Shemer *et al*., 2023). To determine if the ethylene deficiency in the *acs* octuple mutant has been recovered in the spongy mesophyll cell-specific *CORI3::ACS8/11* complementation lines, we measured ethylene production of WT, *acs* octuple, and the three *CORI3::ACS8/11* transgenic lines (#14-14, #16-1, and #31-1). The results revealed a significantly increased ethylene production in lines #14-14 and #16-1 by 2.7 and 3.4 folds when compared to the *acs* octuple mutant plants (Fig. 1 A-4 & S1 A-3; *P*=0.001), while line #31-1 showed an increase by 1.4 folds (Fig. S1 A-3; NS).

#### Stomatal conductance responses to [CO_2_] -shifts

To examine whether complementation of the *acs* octuple mutant, specifically in the spongy mesophyll cells, can restore its impaired stomatal conductance phenotype, we measured stomatal conductance responses to [CO_2_] -shifts in intact leaves of WT, *acs* octuple, and the three *CORI3::ACS8/11* transgenic lines. All three transgenic lines (#14-14, #16-1, and #31-1) displayed similar stomatal conductance values as the *acs* octuple mutant (Fig. 1 B-1 & S1 B-1/C-1), resulted in a similar impaired stomatal conductance response to [CO_2_], (closing slope of WT/*acs* octuple/*CORI3::ACS8/11* = -0.06/-0.015/-0.018; opening slope = 0.04/0.017/0.019) (Fig. 1 B-2&3 for line #14-14; S1 B&C-2&3 for lines #16-1 and #31-1).

#### CO_2_ assimilation rate (*A*) and intrinsic water use efficiency (iWUE)

We found that there was no effect of the *ACS8/ACS11* complementation in the spongy mesophyll on the *A* rates in all three transgenic lines (#14-14, #16-1, and #31-1), which were similar to those of the *acs* octuple (Fig. 1 C-1&2; S1-D&E 1&2). Compatible with its enhanced stomatal closing response, the WT shows a 34.7% higher iWUE rate (*P*=0.007) than the octuple mutant. Notably, there were no significant differences in the iWUE between the transgenic lines and the *acs* octuple mutant (Fig. 1 C-3&4). Similar results were observed among all three transgenic lines measured (S1-D&E 3&4).

### Complementation of *ACS8* and *ACS11*, under both spongy and palisade mesophyll promoters, partially reversed the impaired stomatal conductance responses to [CO_2_]

The complementation of *ACS8* and *ACS11* only in the spongy mesohyl cells of the *acs* octuple mutant did not restore its defective plant phenotype, nor its impaired stomatal conductance responses to [CO_2_] (Fig. 1&S1). To further investigate the role ethylene biosynthesis plays in stomatal response, we generated transgenic Arabidopsis plants that complement the *acs* octuple mutant in both the spongy and palisade mesophyll cells. These lines express the *ACS8* and *ACS11* genes under two promoters, the spongy mesophyll-specific promoter *CORI3* (Procko et al., 2022) and the palisade mesophyll-specific promoter *PEG1* (Geisler *et al*., 2002). Following PCR-based screening (using the binary vector’s primers) for positive lines, three independent *CORI3/PEG1::ACS8/11* lines (e.g., #6-2, #7-1, and #5-6) were selected for detailed physiological and molecular investigations.

#### Plant phenotype

Five-week-old Arabidopsis plants of WT, *acs* octuple, and three different *CORI3/PEG1::ACS8/11* transgenic lines were examined for their anatomical phenotype. All three *CORI3/ PEG1::ACS8/11* transgenic lines conserved the defective leaf phenotype of the *acs* octuple mutant (thinner leaf blade and a downward curling tip) (Fig. 2 A-1&S 2 A-1).

#### Confocal microscopy

*CORI3* and *PEG1* promoters mediate transcription specifically in the spongy and palisade mesophyll, respectively. To evaluate the tissue-specific expression under the *CORI3* and *PEG1* promoters in the transgenic lines, we include in the *CORI3/PEG1::ACS8/11* plasmids also an additional cassette encoding the NEON green fluorescent reporter protein under the *CORI3*/*PEG1* promoter, respectively (provided in Methods S1). Confocal analysis of the leaf cross-section of the *CORI3/PEG1::ACS8/11* transgenic lines revealed that NEON green fluorescent reporter protein expression was limited to the spongy and palisade cells (Fig. 2 A-2).

#### *ACS8* and *ACS11* expression in the leaf by qRT-PCR

To further validate the complementation of the *acs* octuple mutant in the *CORI3/PEG1:: ACS8/11* transgenic lines, RNA was extracted from leaves of 5-weeks-old WT, *acs* octuple, and the three *CORI3/PEG1:: ACS8/11* transgenic lines. qRT-PCR analysis showed low expression levels of *ACS8* and *ACS11* genes in the *acs* octuple mutant compared to their expression levels in WT plants. Fig. 2 A-3 shows the relative gene expression level mean difference ± SEM and *P-*value of the effect size (*P*_ES_): ***ACS8*** -0.4± 0.2; *P*_ES WT – *acs*_ = 0.21; ***ACS11*** - 0.5± 0.04; *P*_ES WT – *acs*_ = 0.008. qRT-PCR analyses of the three transgenic lines (#6-2, #7-1, and #5-6) revealed significantly high expression levels of both *ACS8* and *ACS11* compared to their expression levels in the *acs* octuple mutant. Fig. 2 A-3 shows the mean difference ± SEM and *P-*value of the effect size (*P*_ES_) of line #6-2: ***ACS8*** -65.7±30.4; *P*_ES #6-2 – *acs*_ = 0.16; ***ACS11*** -22.0±12.1, *P*_ES #6-2 – *acs*_ = 0.21. Supplemental table S5 presents raw data for the different *CORI3/PEG1:: ACS8/11*transgenic lines.

#### Ethylene production

To determine whether the deficiency in ethylene production in the *acs* octuple mutant has been recovered in the spongy/palisade mesophyll cell-specific *CORI3/PEG1::ACS8/11* complementation lines, we measured ethylene production of WT, *acs* octuple, and the three *CORI3/PEG1::ACS8/11* transgenic lines. Our analyses showed the highest ethylene production levels in line #6-2, which was not statistically different from ethylene production levels in WT plants and reached 2.7 fold (*P*=0.0015) of its levels in the *acs* octuple mutant (Fig. 2 A-4). Line #7-1 also showed significantly high ethylene production levels, which reached 1.9 fold (*P*=0.0678) of the *acs* octuple mutant production levels. Analysis of the ethylene production level in line #7-1 revealed low ethylene production levels, significantly lower than WT (Fig. S2 A-3). Interestingly, although qRT-PCR showed significantly high transcription levels of *ACS8* and *ACS11* in lines #5-6, ethylene production was slightly higher (NS) than in the *acs* octuple mutant (Fig. S2 A-3).

#### Stomatal conductance responses to [CO_2_] -shifts

We measured stomatal conductance responses to [CO_2_]-shifts in intact leaves of WT, *acs* octuple, and the three *CORI3/PEG1::ACS8/11* transgenic lines. Stomatal conductance steady-state value of the *CORI3/PEG1::ACS8/11* line #6-2 showed a WT like values under 415ppm [CO_2_], which were slightly higher than those of the *acs* octuple mutant (*P* =0.159) (Fig. 2 B-1). As found previously, the *acs* octuple mutant showed an impaired stomatal opening and closing in response to [CO_2_]-shifts. Gas exchange analysis of the *CORI3/PEG1::ACS8/11* line #6-2 showed partial (A 95% confidence interval) complementation of the impaired stomatal opening and closing defect of the *acs* octuple mutant (Fig. 2 B-2&3). The two additional lines measured (#5-6 and #7-1) were comparable to the *acs* octuple mutant stomatal conductance response phenotype (Fig. S2 B&C).

#### CO_2_ assimilation rate (*A*) and intrinsic water use efficiency (iWUE)

The analysis of the *A* rates revealed no significant differences between all three lines (Fig. 2 C-1&2). Analysis of iWUE revealed significantly lower levels of *acs* octuple mutant under high [CO_2_] (900ppm). Interestingly, *CORI3/PEG1*::*ACS8/11* line #6-2 iWUE was similar to WT under all [CO_2_] levels measured and significantly different from the *acs* octuple. This phenotype was not observed in lines #5-6 nor #7-1.

### Complementation of *ACS8* and *ACS11*, specifically in the guard cells of the *acs* octuple mutant, restored the impaired stomatal conductance responses to [CO_2_]-shifts of the *acs* octuple mutant

To test whether complementation of the *acs* octuple mutant specifically in the guard cells can rescue the impaired stomatal conductance response to [CO_2_]-shifts, we generated the *GCD1::ACS8/11* transgenic Arabidopsis line. This line complements the *acs* octuple mutant in a tissue-specific matter expressing the *ACS8 and ACS11* genes under the guard cell-specific promoter GC1 (GCD1) (Yang *et al*., 2008). Three independent *GCD1::ACS8/11* transgenic lines (#8-5, #10-1, and #6-2) were selected based on PCR using the binary vector’s primers (as described in method section 3.6) for in-depth physiological and molecular investigations.

#### *GCD1::ACS8/11* transgenic plants phenotype

5-week-old Arabidopsis plants of WT, *acs* octuple, and three different *GCD1::ACS8/11* transgenic lines were examined for their anatomical plant phenotype. The results show that all three *GCD1::ACS8/11* transgenic lines showed a reverted leaf shape phenotype and standard plant size, similar to the WT plants (Fig. 3A-1 & S9A-1).

#### Confocal microscopy

It has been demonstrated that the GCD1 promoter controls the expression of genes, specifically within the guard cell (Yang *et al*., 2008). To verify that the *GCD1::ACS8/11* construct induces expression specifically in the guard cells, confocal imaging analysis was conducted on leaves from the *acs* octuple mutant (negative control) and the *GCD1::ACS8/11* transgenic lines. Results show that all three *GCD1::ACS8/11* transgenic lines express the NEON reporter protein specifically within the guard cells (Fig. 3 A-2 & S3 A-2).

#### *ACS8* and *ACS11* expression in the guard cells by qRT-PCR

To further validate the complementation of the *acs* octuple mutant within the guard cells, WT, *acs* octuple, and the three *GCD1::ACS8/11* transgenic lines were grown under ambient CO_2_ levels for five weeks. Leaf epidermal peels were extracted, and RNA was isolated. qRT-PCR quantification showed significantly lower expression levels of *ACS8* and *ACS11* genes in the *acs* octuple mutant guard cells when compared to their expression levels in guard cells of WT plants (Fig. 3A-3 shows the mean difference±SEM and *P-value* of the effect size (*P*_ES_): ***ACS8*** -0.53± 0.13; *P*_ES WT – *acs*_ = 0.02; ***ACS11*** -0.48± 0.18; *P*_ES WT – *acs*_ = 0.07). Furthermore, both *ACS8* and *ACS11* were pronouncedly expressed within the guard cells of the three transgenic lines, with line #8-5 displaying the highest gene expression levels when compared to their expression levels in the *acs* octuple mutant guard cells (Fig. 3A-3 shows the relative gene expression level mean difference ± SEM and *P-value* of the effect size (*P*_ES_) of line #8-5: ***ACS8*** ∼529.6± 78.15; *P*_ES #8-5 – *acs*_ = 0.02; ***ACS11*** ∼2136± 589.9, *P*_ES #8-5 – *acs*_ = 0.04). Supplemental table S6 presents raw data for the different *GCD1::ACS8/11* transgenic lines.

The residual expression of *ACS8* and *ACS11* observed in the guard cells of the *acs* octuple mutant is consistent with the genetic strategy used for its construction. While seven members of the *ACS* family were fully inactivated via T-DNA insertions, *ACS8* and *ACS11* were suppressed using artificial microRNA (amiR) technology. Because microRNA-mediated knockdown often results in incomplete silencing rather than a complete knockout, low but measurable transcript levels can persist. This phenomenon was previously noted by Tsuchisaka *et al*. (2009), who reported detectable, albeit significantly reduced, *ACS8* and *ACS11* expression in 5-day-old seedlings of this mutant line.

#### Ethylene production of *GCD1::ACS8/11* transgenic plants

To determine if the ethylene deficiency in the *acs* octuple mutant has been recovered in the guard cell-specific *GCD1::ACS8/11* complementation lines, WT, and the three *GCD1::ACS8/11* transgenic lines (#8-5, #10-1, and #6-2) were measured for their ethylene production using gas chromatography. Results showed a dramatic increase in ethylene production in all three transgenic lines compared to the *acs* octuple mutant (Fig. 3A-4 & S3A-4). Line #8-5 had recovered its ethylene production and reached WT ethylene levels, four times higher than in the *acs* octuple mutant plant leaves (*P*=0.001).

#### Stomatal conductance responses to [CO_2_]-shifts

To examine whether complementation of the *acs* octuple mutant, specifically in the guard-cells, can restore its impaired stomatal conductance phenotype, we measured stomatal conductance responses to [CO_2_]-shifts in intact leaves of WT, *acs* octuple, and the three *GCD1::ACS8/11* transgenic lines. The findings indicated that all three *GCD1::ACS8/11* transgenic lines exhibited higher steady-state levels of stomatal conductance at 415 ppm [CO_2_], comparable to the values observed in the *acs* octuple mutant. In Fig. 3 B-1, the *GCD1::ACS8/11* transgenic line (#8-5) demonstrates an increase in its stomatal conductance steady-state level compared to the acs octuple mutant (P=0.1). Normalized stomatal conductance analysis demonstrated different degrees of recovery of the *acs* octuple mutant impaired stomatal conductance response in the different *GCD1::ACS8/11* transgenic lines (Fig. 3 B-2, #8-5; S3 B-2 and C-2, #10-1 and #6-2). Line #8-5 showed partial stomatal conductance responses to [CO_2_]. In line #8-5, stomatal closure in response to high [CO_2_] increased by 47%, and low [CO_2_]-induced stomatal opening increased by 27% compared to the *acs* octuple mutant (Fig. 3 B-3; *P*<0.05, 95% confidence intervals). Analysis of lines #10-1 showed a smaller effect (Fig. S3 B-3), while line #6-2 showed complete recovery of the *acs* octuple mutant impaired stomatal conductance responses to [CO_2_]-shifts (Fig. S3 C-3; *P*<0.05, 95% CI).

#### CO_2_ assimilation rate (*A*) and intrinsic water use efficiency (iWUE)

Compatible with its high stomatal conductance values at 415ppm [CO_2_], the *GCD1::ACS8/11* transgenic line exhibited higher *A* rates and lower iWUE compared to the WT (*P*=0.019 and *P*=0.061, respectively) (Fig. 3 C-1&2). The *acs* octuple and *GCD1::ACS8/11* also showed a higher *A* rate at 900ppm (*P*=0.063) with higher iWUE of (*P*=0.037 and *P*=0.063 respectively) (Fig. 3 C-1&2).

### Complementation of *ACS8* and *ACS11* under the whole-plant expression promoter resulted in dwarf sterile plants

To investigate whether complementation of the *ACS8* and *ACS11* genes in the *acs* octuple mutant under a constitutive UBQ10 promoter may repair the impaired stomatal conductance response to [CO_2_]-shifts, we generated *UBQ10::ACS8/11* transgenic Arabidopsis lines. Many attempts were made to create positive and vital transgenic plants, yet our attempts resulted only in two lines showing a severe dwarf and sterile phenotype (Fig. S4). As these plants were weak and didn’t survive, no additional measurements could be performed on these lines.

## Discussion

Ethylene, a gaseous phytohormone, regulates stomatal conductance in response to [CO_2_], coordinating stomatal conductance with the plant’s overall gas exchange needs (Desikan *et al*., 2006; Fujita *et al*., 2013). In an earlier study, [CO_2_] was found to induce ethylene production in Arabidopsis leaves and mediate stomatal conductance (Azoulay-Shemer *et al*., 2023). Different studies have pointed out a complex regulation of ethylene biosynthesis in a tissue-dependent matter. As ethylene may affect stomata directly via guard-cell ethylene biosynthesis and perception (Desikan *et al*., 2006; Beguerisse-Diaz *et al*., 2012), and/or indirectly as long-distance mesophyll-driven signals (Lee & Bowling, 1995; Fujita *et al*., 2013; Azoulay-Shemer *et al*., 2023), we aim to elucidate the ethylene-producing tissue/s that are involved in [CO_2_]-induced stomatal conductance regulation.

In this study, we generated four tissue-specific complementation lines, complementing the *acs* octuple mutant using four different tissue-specific promoters for the spongy mesophyll, spongy, and palisade mesophyll, guard cell, or whole plant. Our study revealed that guard cell-specific complementation was most effective in restoring the *acs* octuple mutant impaired stomatal conductance responses to [CO_2_], while mesophyll-specific complementation only partially rescued these phenotype.

### Tissue-specific ethylene biosynthesis and its effect on stomatal conductance responses to CO_2_

Ethylene biosynthesis is governed by a complex, multilevel control circuitry (Pattyn *et al*., 2021) and is tightly regulated through diverse mechanisms that vary according to tissue type (Wang *et al*., 2022). The results of this study indicate that both mesophyll and guard cell ethylene biosynthesis contribute to stomatal conductance regulation, with guard cell-derived ethylene playing a primary role. While mesophyll-produced ethylene complementation lines (*CORI3*/*PEG1*::*ACS8/11*) elevates overall leaf ethylene levels, it appears insufficient for optimal stomatal function. Nevertheless the partial (significant) rescue of stomatal conductance responses to [CO_2_] indicate that mesophyll-derived ethylene may play a supportive role in stomatal regulation, possibly through long-distance signaling or by contributing to the overall ethylene pool in the leaf. This finding is consistent with the idea of mesophyll-to-epidermis signaling in stomatal regulation (Mott *et al*., 2008; Azoulay-Shemer *et al*., 2023).

#### Mesophyll

The results of this study showed that whole mesophyll-specific complementation lines (spongy&palisade; *CORI3*/*PEG1*::*ACS8/11*), but not the spongy mesophyl-specific complementation lines (*CORI3*::*ACS8/11*), parially restored stomatal CO₂ responses. The leaf mesophyll constitutes the bulk of the leaf volume and is differentiated into specialized palisade and spongy mesophyll layers. The spongy mesophyll, located close to the stomatal pores on the abaxial side of the leaf, facilitates gas exchange. As these cells are the most exposed tissue in the leaf to [CO_2_] gas exchange, they may be involved in CO_2_-sensing and regulation of stomatal conductance using ethylene as long-distance signaling. Interestingly, complementation of the *acs* octuple mutant in the spongy mesophyll cells alone (*CORI3*::*ACS8/11*) didn’t restore the impaired stomatal conductance responses to [CO_2_] (Fig. 2B). Palisade cells, by contrast, constitute the primary photosynthetic tissue of the leaf, optimized for maximum light absorption and carbon fixation. Various studies showed that photosynthesis products, such as sugar, mediate stomatal conductance to optimize leaf gas exchange and the plant’s water use efficiency (Lee & Bowling, 1995; Baroli *et al*., 2008; Kelly *et al*., 2019). Given these considerations, mesophyll tissues are likely to contribute to the regulation of stomatal conductance, potentially using gaseous signals such as ethylene as part of the mesophyll-to-guard-cell communication network (Sobeih *et al*., 2004; Azoulay-Shemer *et al*., 2023). Remarkably, when the *acs* octuple mutant was complemented in both the spongy and palisade mesophyll (*CORI3/PEG1*::*ACS8/11*), although it produced ethylene to the same levels as in the spongy mesophyll complementation line, it led to a partial (significant) rescue of stomatal conductance responses to [CO_2_] (Fig. 2 B).

#### Guard cells

The results of this study showed that guard-cell-specific complementation lines (*GCD1::ACS8/11*) most effectively restored stomatal CO₂ responses, yet not fully. Guard cells have been previously shown to respond to various signals autonomously (Kim *et al*., 2010; Zhang *et al*., 2021). Different studies have demonstrated that guard cells express the ethylene biosynthesis pathway key enzymes, including ACC synthase and ACC oxidase (Merritt *et al*., 2001; Yin *et al*., 2019), and can synthesize ethylene autonomously (Beguerisse-Diaz *et al*., 2012). Furthermore, ethylene was found to be perceived by guard cells in isolated epidermal peels (Levitt *et al*., 1987; Tanaka *et al*., 2005) and triggers various pathways involved in the regulation of stomatal conductance (Desikan *et al*., 2004), including the synthesis of hydrogen peroxide (Desikan *et al*., 2006; Watkins *et al*., 2014). However, based on a detailed study that showed that ethylene induces stomatal closure in leaves but not in epidermal peels, Desikan *et al*. (2006) suggested that ethylene-induced stomatal closure requires communication between guard cells and mesophyll cells.

Ethylene production, perception, and signaling mechanisms operate in guard cells (Tanaka *et al*., 2005; Beguerisse-Diaz *et al*., 2012; Munemasa *et al*., 2019). ACS8 expression shows robust circadian oscillations that closely track ethylene emission in Arabidopsis, indicating that ACC SYNTHASE 8 (ACS8) is a major clock-regulated control point for ethylene production under the control of CCA1/TOC1 (Thain *et al*., 2004). Moreover, ethylene signaling can feed back on the circadian oscillator, with ACC treatments and CTR1-dependent pathways reported to modulate circadian period and sustain clock reporter rhythms (Haydon *et al*., 2017). If the GCD1::ACS8/11 genetic modifications alter circadian regulation of ethylene biosynthesis specifically in guard cells, this could shift the phase or amplitude of guard-cell clock outputs and thereby modify stomatal behavior across the photoperiod (Haydon *et al*., 2017; Yari Kamrani *et al*., 2022), which is consistent with the elevated stomatal conductance observed in these lines (Fig. 3B).

Ethylene plays a crucial role in regulating stomatal conductance, in part by triggering H₂O₂-dependent signaling in guard cells and engaging Ca²⁺- and anion-channel-mediated pathways that can lead to stomatal closure (Desikan *et al*., 2006; Munemasa *et al*., 2019). Nevertheless, ethylene’s effects are context dependent and can either promote or inhibit stomatal closure through cross-talk with other signaling pathways and guard-cell ion channels, rather than acting as a simple opener or closer (Munemasa *et al*., 2019). CO₂ also regulates stomatal conductance by activating S-type anion channels and outward-rectifying K⁺ channels and, via the resulting depolarization, reducing inward-rectifying K⁺ currents, changes that are associated with stomatal closure (Brearley *et al*., 1997; Schroeder *et al*., 2001a). In addition, elevated CO₂ has been shown to modulate cytosolic Ca²⁺ dynamics in guard cells, and CO₂-induced Ca²⁺ elevations are implicated in stomatal responses, suggesting that CO₂-regulated Ca²⁺ signaling forms an integral component of the stomatal CO₂ response (Webb *et al*., 1996). In this study, guard-cell-specific complementation lines (GCD1::ACS8/11) most effectively restored stomatal CO₂ responses, consistent with a key role for localized ethylene biosynthesis within guard cells in stomatal regulation. We therefore hypothesize that guard-cell-localized ethylene production influences ion channel activity, in concert with CO₂ signaling, to fine-tune stomatal conductance responses to changes in [CO₂].

### Spatial Regulation of the Ethylene Biosynthetic Pool

A significant finding of our study was the inability to recover viable plants expressing ACS8 and ACS11 under the constitutive UBQ10 promoter, which underscores the critical importance of spatial specificity in ethylene signaling. Unlike our tissue-specific lines, the UBQ10::ACS8/11 lines exhibited a severe dwarf and sterile phenotype (Fig. S4), consistent with high, unregulated ethylene production throughout the plant. These results highlight the challenges of manipulating hormone signaling, where loss of fine-tuned, localized expression can lead to detrimental phenotypes. Similar strong growth inhibition and developmental defects have been described in plants with constitutively activated ethylene responses (Kieber *et al*., 1993; Pierik *et al*., 2006). This outcome further emphasizes that the spatial separation of ethylene biosynthesis—rather than a simple global increase in the hormone pool—is essential for balancing normal plant development with specialized physiological responses, such as stomatal regulation

### Ethylene, CO_2_ assimilation, and water use efficiency

Analysis of CO₂ assimilation rates (*A*) and intrinsic water-use efficiency (iWUE) across the different *acs* octuple mutant tissue-specific complementation lines revealed intriguing patterns. Previous studies have shown that ethylene can affect stomatal conductance (Tanaka *et al*., 2005; Azoulay-Shemer *et al*., 2023) and modulate photosynthetic capacity by influencing chlorophyll biosynthesis, chlorophyll content, and the performance of photosynthetic complexes (Alscher & Castelfranco, 1972; Ceusters & Van de Poel, 2018). In this study, the *acs* octuple mutant, which is impaired in ethylene production, showed similar *A* rates as WT plants under all [CO_2_] measured, while its impaired stomatal closure under high CO_2_ levels resulted in reduced iWUE, indicating that the *acs* octuple mutant is less efficient in balancing carbon gain with water loss under these conditions. Interestingly, our investigation reveals that the complementation of the *acs* octuple mutant, specifically within its guard cells (*GCD1::ACS8/11*) lines, but not in the spongy or palisade mesophyll, resulted in higher *A* rates under ambient CO_2_ levels (415ppm). This result is in line with the higher gs steady state level that was ditected solely in the guard cells (*GCD1::ACS8/11*) lines. Furthermore, it has been shown that ethylene involve in stomatal development (Serna & Fenoll, 1996; Serna & Fenoll, 2000; Saibo *et al*., 2003; Gong *et al*., 2021). Indeed our data showed that guard cells specific complementation of the *acs* mutant can recover the impaired stomatal development of the *acs* octuple mutant (i.e., lower stomatal indices, reduced stomatal density, and a larger stomatal area when compeared to WT) (Data not shown). Facinetly, while the mesophyll-specific (palisade and spongy) complementation line *CORI3/PEG1*::*ACS8/11* displayed iWUE rates comparable to the WT, the spongy cells specific complementation lines (*CORI3*::*ACS8/11*) and guard cells specific (*GCD1::ACS8/11*) complementation lines demonstrated comparable iWUE levels as the *acs* octuple mutant. The results highlights the complex interplay between ethylene production sites and their physiological effects, emphasizing the importance of spatially regulated ethylene biosynthesis in balancing gas exchange, CO_2_ assimilation and intrinsic water-use efficiency (iWUE).

In summary, our investigation presents strong evidence for the pivotal role of localized ethylene biosynthesis and its role in guard cells and mesophyll cells in modulating stomatal response in Arabidopsis thaliana. These findings underscore the intricate nature of ethylene signaling in plant physiology and suggest novel approaches for crop enhancement through cell-type-specific manipulation of ethylene biosynthesis.

## Acknowledgements

We thank Sakis Theologis (Albany, California) for providing the ACC-synthase octuple mutant. We thank Prof. Julian Schroeder (UCSD, California) for providing the guard cell-specific promoter GCD1 construct, for scientific feedback and discussions. We thank Dr. Carl Procko and Prof. Joanne Chory (Salk Ins, CA, USA) for providing the CORI3 and PEG1-specific promoter constructs. This research was funded by the Israel Science Foundation (grant no. 768/20) to TA-S.

## Author contributions

TA-S conceptualized and designed this research. DN-R planned and performed all molecular biology and the generation of transgenic plants. DN-R, SO and DN designed and performed phenotyping and physiological analyses. DN-R conducted ethylene production analyses and qPCR.

DN-R and GS performed all microscopic analyses.

## Data availability

The data that support the findings of this study are available in the Supporting Information (Figuers S1-S4, Tables S1-S6 and Method S1) of this article.

## Supporting Information

The following Supporting Information is available for this article:

**Fig. S1 Physiological and molecular characterization of *CORI3::ACS8/11* transgenic *lines (#16-1 and #31-1.)***

**Fig. S2 Physiological and molecular characterization of *CORI3/PEG1::ACS8/11* transgenic *lines* (#5-6 and #7-1).**

**Fig. S3 Physiological and molecular characterization of *GCD1::ACS8/11* transgenic *lines***

**Fig. S4 Characterization of *UBQ10::ACS8/11* transgenic *line*.**

**Table S1 List of PCR oligonucleotide primers used for the plasmids construction.**

**Table S2 List of PCR oligonucleotide primers used for transgenic line verification.**

**Table S3 List of qRT-PCR oligonucleotide primers.**

**Table S4 qRT-PCR of *ACS*8 and *ACS11* genes in the *CORI3::ACS8/11* transgenic lines.**

**Table S5 qRT-PCR of *ACS*8 and *ACS11* genes in the *CORI3/PEG1::ACS8/11* transgenic lines.**

**Table S6 qRT-PCR of *ACS*8 and *ACS11* genes in the *GCD1::ACS8/11* transgenic lines.**

**Methods S1 Schematics of DNA constructs and expression promoters.**

