## Supplementary material for "Ethylene biosynthesis in guard cells, not mesophyll, predominantly drives stomatal conductance responses to CO_2_": Supp files

The following Supporting Information is available for this article:

**Fig. S1 Physiological and molecular characterization of *CORI3::ACS8/11* transgenic lines *(#16-1 and #31-1.)***

**Fig. S2 Physiological and molecular characterization of *CORI3/PEG1::ACS8/11* transgenic lines (#5-6 and #7-1).**

**Fig. S3 Physiological and molecular characterization of *GCD1::ACS8/11* transgenic lines**

**Fig. S4 Characterization of *UBQ10::ACS8/11* transgenic line.**

**Table S1 List of PCR oligonucleotide primers used for the plasmids construction.**

**Table S2 List of PCR oligonucleotide primers used for transgenic line verification.**

**Table S3 List of qRT-PCR oligonucleotide primers.**

**Table S4 qRT-PCR of *ACS*8 and *ACS11* genes in the *CORI3::ACS8/11* transgenic lines.**

**Table S5 qRT-PCR of *ACS*8 and *ACS11* genes in the *CORI3/PEG1::ACS8/11* transgenic lines.**

**Table S6 qRT-PCR of *ACS*8 and *ACS11* genes in the *GCD1::ACS8/11* transgenic lines.**

**Methods S1 Schematics of DNA constructs and expression promoters.**

**Supplemental Figuers**

**Fig. S1 Physiological and molecular characterization of *CORI3::ACS8/11* transgenic *lines (#16-1 and #31-1).*** 5-week-old WT (Col-0), the *acs* octuple mutant, and the *CORI3::ACS8/11* transgenic Arabidopsis plants (lines *#16-1 and #31-1)* express the *ACS*8 *and ACS*11 under the spongy mesophyll cell-specific promoter CORI3. All lines were analyzed for the following measurements: **(A) 1)** Plant phenotype **2) qRT-PCR of** *ACS*8 and *ACS*11 genes expression in leaf samples. Data were normalized to the *ACT8* expression level. Results are presented as relative gene expression levels ($log^2$) ±SEM (n=3) plants per genotype; $P<0.05$, two-way ANOVA followed by Fisher's LSD test. **3)** Ethylene production levels of *whole plant* rosettes ±SEM (WT (n=6), *acs* octuple (n=6), line #16-1 (n=3) and line #31-1 (n=7); $P<0.05$, one-way ANOVA followed by Dunnett test). **(B &C)** Time-resolved stomatal conductance responses were analyzed at the imposed $CO_2$ shifts (indicated at the top in ppm) in intact plant leaves of WT, acs octuple and *CORI3::ACS8/11*. **(B)** line #16-1 (**C**) line #31-1. 1) Stomatal conductance in mol $H_2O$ $m^{-2}$ $sec^{-1}$ 2) Relative stomatal conductance. Data were normalized to the stomatal conductance at the last 15-sec time-point under 415 ppm [$CO_2$] before [$CO_2$] was shifted to 900 ppm. 3) Relative stomata closing and opening rates were calculated with a linear regression line (±SEM). The line was calculated between the $5^{th}$ and $20^{th}$ minute, under high or low [$CO_2$], respectively, and the linear least squares regression statistical fitting method was used. A 95% confidence interval was used for means comparison within each line. **(D&E)** 1) Net $CO_2$-assimilation rates (µmol $CO_2$ $m^{-2}$ $sec^{-1}$) **(D)** line #16-1 **(E)** line #31-1 (2) The averaged net assimilation rate was calculated as the mean of the final data point for each $CO_2$ level (i.e., 415, 900, 100ppm). (3) Intrinsic water use efficiency (iWUE). (4) Averaged intrinsic water use efficiency was calculated as the mean of the final data point for each $CO_2$ level. Data are the mean (±SEM) of n=5 leaves from individual plants per genotype. *P*-value style was mentioned as its numeric value.

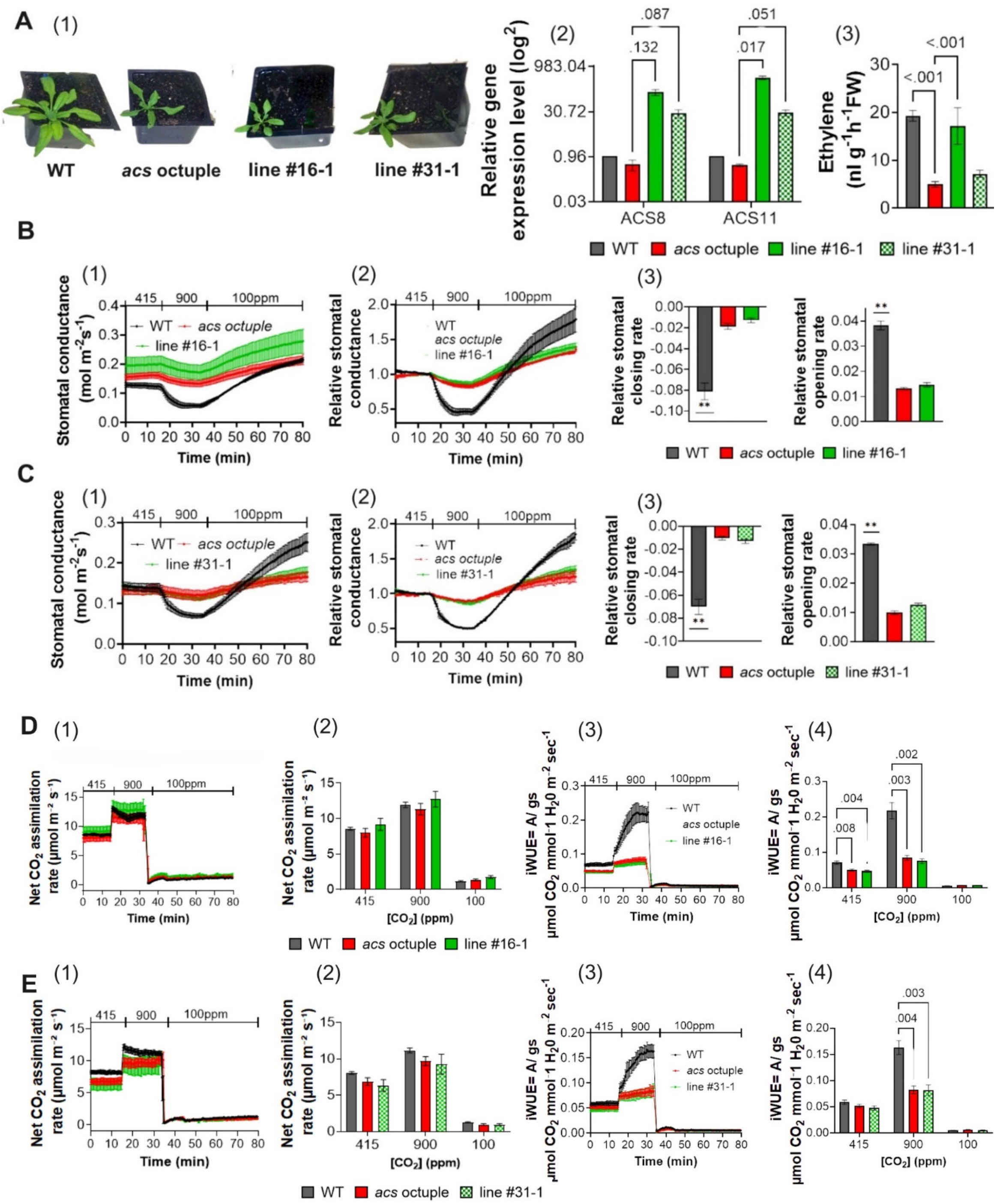

A (1)
WT
acs octuple
line #16-1
line #31-1
(2)
Relative gene expression level (log2)
983.04
30.72
0.96
0.03
.087
.132
.051
.017
ACS8
ACS11
(3)
Ethylene (nl g-1h-1FW)
<.001
<.001
WT
acs octuple
line #16-1
line #31-1
B (1)
Stomatal conductance (mol m-2s-1)
415
900
100ppm
Time (min)
(2)
Relative stomatal conductance
(3)
Relative stomatal closing rate
Relative stomatal opening rate
**
C (1)
(2)
(3)
D (1)
Net CO2 assimilation rate (µmol m-2 s-1)
(2)
[CO2] (ppm)
(3)
iWUE= A/ gs
µmol CO2 mmol-1 H20 m-2 sec-1
(4)
.008
.004
.003
.002
E (1)
(2)
(3)
(4)
.004
.003

**Fig. S2 Physiological and molecular characterization of *CORI3/PEG1::ACS8/11* transgenic *lines* (#5-6 and #7-1).** 5-week-old WT, *acs* octuple mutant, and the *CORI3/PEG1::ACS8/11* transgenic Arabidopsis plants (line #6-5 and #7-1) expressing the *ACS8* and *ACS11* under the spongy mesophyll cell-specific promoter, *CORI3*, and the palisade mesophyll, *PEG1*. All lines were analyzed for the following measurements: **(A) 1)** Plant phenotype **2) qRT-PCR of** *ACS*8 and *ACS*11 genes expression in leaf samples. Data were normalized to the *ACT8* expression level. Results are presented as relative gene expression levels ($log^2$) ±SEM (n=3) plants per genotype; $P<0.05$, two-way ANOVA followed by Fisher's LSD test. **3)** Ethylene production levels of whole plant rosettes ±SEM (WT (n=6), *acs* octuple (n=6), lines #5-6 and #7-1; $P<0.05$, one-way ANOVA followed by Dunnett test). **(B &C)** Time-resolved stomatal conductance responses were analyzed at the imposed $CO_2$ shifts (indicated at the top in ppm) in intact plant leaves of WT, acs octuple, and *CORI3/PEG1::ACS8/11*. **(B)** line #6-5 **(C)** line #7-1. 1) Stomatal conductance in mol $H_2O$ $m^{-2}$ $sec^{-1}$ 2) Relative stomatal conductance. Data were normalized to the stomatal conductance at the last 15-sec time-point under 415 ppm [$CO_2$] before [$CO_2$] was shifted to 900 ppm. 3) Relative stomata closing and opening rates were calculated with a linear regression line (±SEM). The line was calculated between the $5^{th}$ and $20^{th}$ minute, under high or low [$CO_2$], respectively, and the linear least squares regression statistical fitting method was used. A 95% confidence interval was used for means comparison within each line. **(D&E)** 1) Net $CO_2$-assimilation rates (µmol $CO_2$ $m^{-2}$ $sec^{-1}$) **(D)** line #5-6 **(E)** line #7-1 (2) The averaged net assimilation rate was calculated as the mean of the final data point for each $CO_2$ level (i.e., 415, 900, 100ppm). (3) Intrinsic water use efficiency (iWUE). (4) Averaged intrinsic water use efficiency was calculated as the mean of the final data point for each $CO_2$ level. Data are the mean (±SEM) of n=5 leaves from individual plants per genotype. *P*-value style was mentioned as its numeric value.

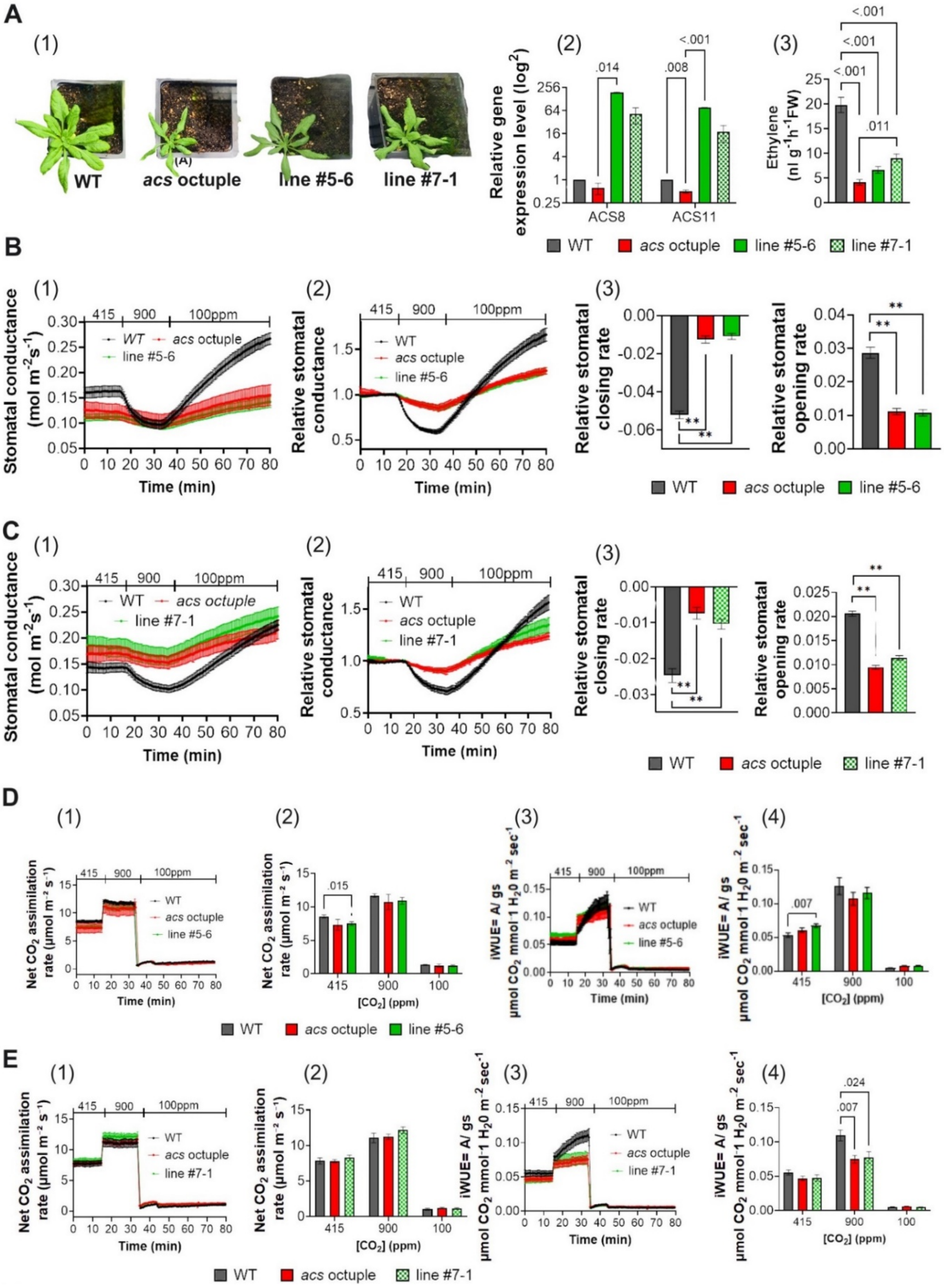
A
(1)
WT
acs octuple
line #5-6
line #7-1
(2)
Relative gene expression level (log2)
ACS8
ACS11
.014
.008
<.001
(3)
Ethylene (nl g-1h-1FW)
<.001
<.001
<.001
.011
WT
acs octuple
line #5-6
line #7-1
B
(1)
415
900
100ppm
Stomatal conductance (mol m-2s-1)
Time (min)
WT
acs octuple
line #5-6
(2)
Relative stomatal conductance
Time (min)
(3)
Relative stomatal closing rate
Relative stomatal opening rate
**
WT
acs octuple
line #5-6
C
(1)
415
900
100ppm
Stomatal conductance (mol m-2s-1)
Time (min)
WT
acs octuple
line #7-1
(2)
Relative stomatal conductance
Time (min)
(3)
Relative stomatal closing rate
Relative stomatal opening rate
**
WT
acs octuple
line #7-1
D
(1)
Net CO2 assimilation rate (µmol m-2 s-1)
Time (min)
(2)
Net CO2 assimilation rate (µmol m-2 s-1)
[CO2] (ppm)
.015
(3)
iWUE= A/ gs
µmol CO2 mmol-1 H20 m-2 sec-1
Time (min)
(4)
iWUE= A/ gs
µmol CO2 mmol-1 H20 m-2 sec-1
[CO2] (ppm)
.007
WT
acs octuple
line #5-6
E
(1)
Net CO2 assimilation rate (µmol m-2 s-1)
Time (min)
(2)
Net CO2 assimilation rate (µmol m-2 s-1)
[CO2] (ppm)
(3)
iWUE= A/ gs
µmol CO2 mmol-1 H20 m-2 sec-1
Time (min)
(4)
iWUE= A/ gs
µmol CO2 mmol-1 H20 m-2 sec-1
[CO2] (ppm)
.024
.007
WT
acs octuple
line #7-1

**Fig. S3 Physiological and molecular characterization of *GCD1::ACS8/11* transgenic *lines.*** 5-week-old WT (Col-0), the *acs* octuple mutant, and the *GCD1::ACS8/11* transgenic Arabidopsis plant (lines *#10-1 and #6-2)* express the *ACS8 and ACS11* under the guard cell-specific promoter GCD1 were analyzed for the following measurements: **(A) 1)** Plant phenotype **2)** Light microscope analysis of a leaf cross-section displays the NEON green fluorescent protein, which is expressed in guard cells nucleus under the GCD1 promoter. The size bar on the bottom left equals 20μm **3) qRT-PCR of** *ACS8* and *ACS11* genes expressed in guard cells from enriched epidermal peel leaf samples. Data were normalized to the WT *ACT8* gene expression level. Results are presented as relative gene expression levels ($log^2$) ±SEM (n=3) plants per genotype; $P<0.05$, two-way ANOVA followed by Fisher's LSD test. **4)** Ethylene production levels of *whole plant* rosettes ±SEM (WT (n=6), *acs* octuple (n=3) , lines #10-1 and #6-2 (n=5); $P<0.05$, one-way ANOVA followed by Dunnett test). **(B&C)** Time-resolved stomatal conductance responses were analyzed at the imposed [$CO_2$]-shifts (indicated at the top in ppm) in intact plant leaves of WT, *acs* octuple and *GCD1::ACS8/11* **(B)** line #10-1 **(C)** line #6-2 1) Stomatal conductance in mol $H_2O$ $m^{-2}$ $sec^{-1}$ 2) Relative stomatal conductance. Data were normalized to the stomatal conductance at the last 15-sec time-point under 415 ppm [$CO_2$] before [$CO_2$] was shifted to 900 ppm. 3) Relative stomata closing and opening rates were calculated with a linear regression line (±SEM). The line was calculated between the $5^{th}$ and $20^{th}$ minute, under high or low [$CO_2$], respectively, and the linear least squares regression statistical fitting method was used. A 95% confidence interval was used for means comparison within each line. **(D&E)** 1) Net $CO_2$-assimilation rates (μmol $CO_2$ $m^{-2}$ $sec^{-1}$) **(D)** line #10-1 **(E)** line #6-2 (2) The averaged net assimilation rate was calculated as the mean of the final data point for each $CO_2$ level (i.e., 415, 900, 100ppm). (3) Intrinsic water use efficiency (iWUE). (4) Averaged intrinsic water use efficiency was calculated as the mean of the final data point for each $CO_2$ level. Data are the mean (±SEM) of n=5 leaves from individual plants per genotype. *P*-value style was mentioned as its numeric value.

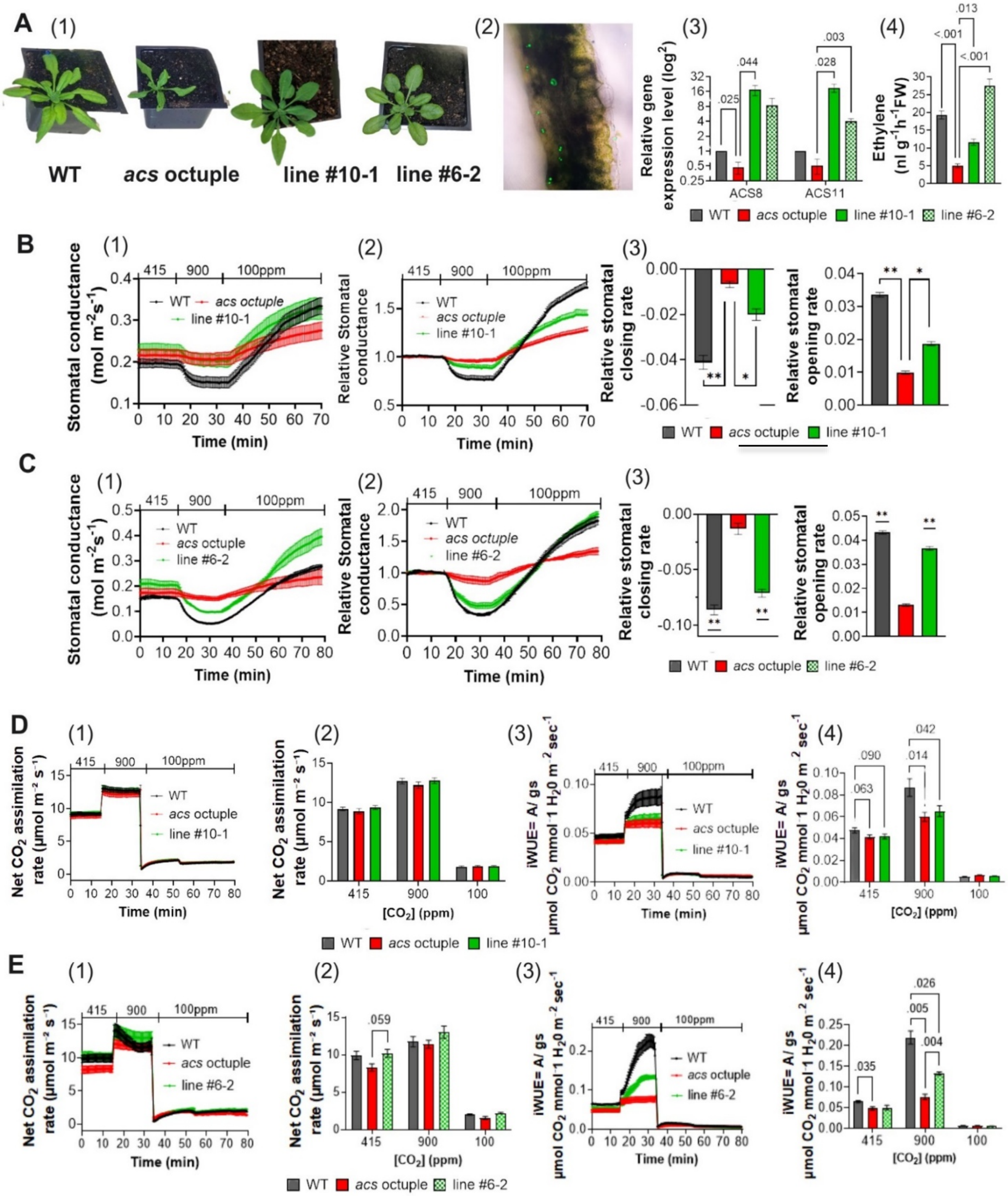
A
(1)
WT
acs octuple
line #10-1
line #6-2
(2)
(3)
Relative gene expression level (log2)
ACS8
ACS11
.025
.044
.028
.003
(4)
Ethylene (nl g-1h-1FW)
<.001
.013
<.001
WT
acs octuple
line #10-1
line #6-2
B
(1)
Stomatal conductance (mol m-2s-1)
Time (min)
415
900
100ppm
(2)
Relative Stomatal conductance
(3)
Relative stomatal closing rate
Relative stomatal opening rate
C
D
Net CO2 assimilation rate (µmol m-2 s-1)
[CO2] (ppm)
iWUE= A/ gs
µmol CO2 mmol 1 H20 m-2 sec-1
.063
.090
.014
.042
E
.059
.035
.005
.026
.004

**Fig. S4 Characterization of *UBQ10::ACS8/11* transgenic *line.*** 5-week-old WT, *acs* octuple mutant, and the *UBQ10::ACS8/11* transgenic Arabidopsis plants (lines *#10),* which express the *ACS*8 *and ACS*11 under the Polyubiquitin10 gene promoter (UBQ10), which promotes high transcription levels in all plant tissues. **(A)** Plant phenotype **(B)** Light microscopy of the NEON green fluorescent protein, detected in the nucleus of most plant cells. The size bar on the bottom left equals 20µm**.**

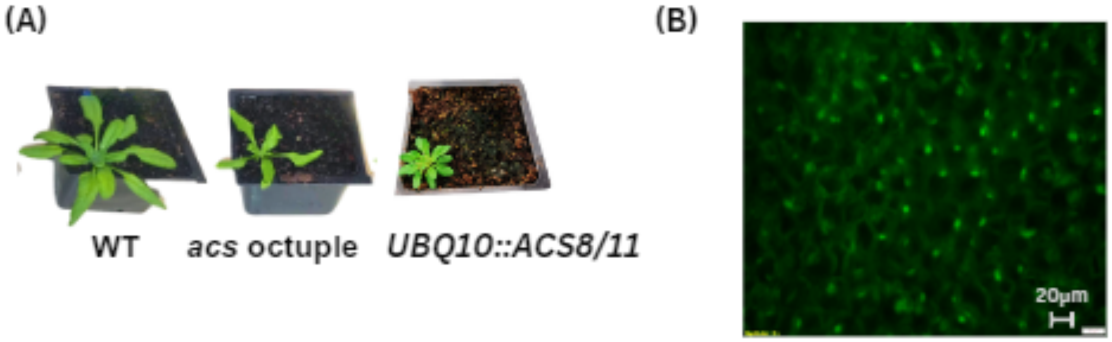

### Supplemental Tables

### Table S1 List of PCR oligonucleotide primers used for the plasmids construction.

| Gene name | Function | Final plasmid constructed name | Primer name | Primer used for L0 level 5'→3' orientation |
|---|---|---|---|---|
| CORi3 (CORI3) | Promoter | pICSL4723 L2 Cori3 promoter + NEON | Cori3 pcr1 F | GAAGACCGGGAGCTCTCTAGATTTTCAGAAGATT |
| | | | Cori3 pcr1 R | GAAGACAATCTCTGTTTTCGACCGACGAG |
| | | | Cori3 pcr2 F | GAAGACAAGAGACAAAAAAATATTTAAATACG |
| | | | Cori3 pcr2 R | GAAGACGGGGATATTAACATCAACATAGCAGACA |
| | | | Cori3 pcr3 F | GAAGACATATCCCACGTGAGATATCA |
| | | | Cori3 pcr3 R | GAAGACTTCATTGACTAACTCTCATTGCTACGAAG |
| GC1 (GCD1) | Promoter | pICSL4723 L2 GCD1 promoter + NEON | GC1 pcr1 F | AAGAAGACTGGGAGATGGTTGCAACAGAGAGGATGAATTTATAAG |
| | | | GC1 pcr1 R | TTGAAGACAACATTCTTGAGTAGTGATTTTGAAGTAGTGTGTGA |
| IQD22 (PEG1) | Promoter | pICSL4723 L2 PEG1 promoter + NEON | PEG1 pcr1 F | AAGAAGACTGGGAGGTAAACAGTACATGGTTAAAGGATAGTTC |
| | | | PEG1 pcr1 R | TTGAAGACAAAGATCCACCAACACAAGCGATTATGG |
| | | | PEG1 pcr2 F | AAGAAGACTGATCTATAGTAATCTAAACAATTTTTTATTTCCATGG |
| | | | PEG1 pcr2 R | TTGAAGACAACATTCTAATGAAAGTTACTTGACGAATGAAACGA |
| UBIQUITIN 10 | Promoter | pICSL4723 L2 UBQ10 promoter + NEON | UBQ10 pro F | AAGAAGACTGGGAGCAGTCTAGCTCAACAGA |
| | | | UBQ10 pro R | TTGAAGACAATTTCTGATTAACAGGTTC |
| ACS8 | CDS | ACS8 L0 | ACS8 CDS F | AAGAAGACTGAATGGGTCTATTGTCAAAGAAAGCTAGTTGC |
| | | | ACS8 CDS R | TAGAAGACAAAAGCCTATCGTTCCTCGGGTTCACGGTCGTGAAA |
| ACS11 | CDS | ACS11 L0 | ACS11 CDS F | TCCCGGGATGTTGTCAAGCAAAGTTGTTGGC |
| | | | ACS11 CDS R | GTTGGATCCTCAACGTTCTGATTCACAAGTAACAGAGG |

### Table S2 List of PCR oligonucleotide primers used for transgenic line verification.

| Primer name | Transgenic line promoter | Plasmid sequence 5'→3' orientation |
|---|---|---|
| Cori3 F | Cori3 | GAAGACAAGAGACAAAAAAATATTTAAATACG |
| RB Rev | Cori3 | GGTTTACCCGCCAATATATCCTGTCA |
| 6288 Venus F | GCD1 | AAAGACCCCAACGAGAAGCGCG |
| 6851 NOS R | GCD1 | AACATAGATGACACCGCGCGCG |
| 7164 peg1 F | PEG1 | GCAATAATATCTCACTATACAGTCC |
| acs11 11904 R | PEG1 | CTGGCAAGCCATGGTAATCT |
| UBQ 5681 F | UBQ10 | TCGTTAAATCTCAACGGCTGGATC |
| 189 acs8 R | UBQ10 | CTCTTTGGAAATTGGCTGCGTCG |

**Table S3 List of qRT-PCR oligonucleotide primers.**

| Primers name | The sequence is shown in 5'→3' orientation | Description |
|---|---|---|
| AtACS8 F | CCTTCCTTCCTTCAAGAATGCTAT | primers for AtACS8 (At4g37770) |
| AtACS8 R | GAGAGTCTCGTTAGCCGGAGTA | |
| AtACS11 F | TGGATGGCAAGAATACGAGAAG | primers for AtACS11 (AT4G08040) |
| AtACS11 R | AACCCAAGACTTCTGGATGC | |
| AtActin8 | CAGCACTTTCCAGCAGATGT | primers for AtAct8 (AT1G49240) |
| AtActin8 | TGATCCCGTCATGGAAACGA | |

**Table S4 qRT-PCR of *ACS8* and *ACS11* genes in the *CORI3::ACS8/11* transgenic lines.** The table shows ACS8 and ACS11 gene expression levels in the acs octuple mutant compared to WT and the CORI3::ACS8/11. The tables present the relative ACS8 and ACS11 gene expression means difference, 95% confidence intervals, and P-value of the effect size (PES) of each separate comparison. The mean difference presents the averaged expression level in the acs octuple mutant compared to each line. The model parameters were analyzed as repeated measurements using two-way ANOVA followed by Fisher's LSD test.

| Table S4 | Mean difference ± SE | 95% CI of diff. | $P_{ES}$ | Mean difference ± SE | 95% CI of diff. | $P_{ES}$ |
|---|---|---|---|---|---|---|
| *CORI3::ACS8/11* | ***ACS8*** | | | ***ACS11*** | | |
| *acs* octuple vs. WT | -0.5±0.2 | -1.4 to 0.4 | 0.16 | -0.5±0.06 | -1.3 to 0.3 | 0.08 |
| *acs* octuple vs. line #14-14 | -38.9±26.8 | -154.3 to 76.6 | 0.28 | -17.9±4.5 | -75.1 to 39.2 | 0.15 |
| *acs* octuple vs. line #16-1 | -136.8±28.7 | -501.9 to 228.2 | 0.13 | -414.9±54.1 | -647.7 to -182.0 | 0.01 |
| *acs* octuple vs. line #31-1 | -26.5±8.4 | -62.5 to 9.5 | 0.08 | -28.0±6.6 | -56.3 to 0.3 | 0.05 |

**Table S5 qRT-PCR of *ACS8* and *ACS11* genes in the *CORI3/PEG1::ACS8/11* transgenic lines.** The table shows ACS8 and ACS11 gene expression levels in the acs octuple mutant compared to WT and the CORI3/PEG1::ACS8/11 . The tables present the relative ACS8 and

ACS11 gene expression means difference, 95% confidence intervals, and P-value of the effect size (PES) of each separate comparison. The mean difference presents the averaged expression level in the acs octuple mutant compared to each line. The model parameters were analyzed as repeated measurements using two-way ANOVA followed by Fisher's LSD test.

| **Table S5** | **Mean difference ± SE** | **95% CI of diff.** | $P_{ES}$ | **Mean difference ± SE** | **95% CI of diff.** | $P_{ES}$ |
|---|---|---|---|---|---|---|
| *CORI3/PEG1::ACS8/11* | ***ACS8*** | | | ***ACS11*** | | |
| *acs* octuple vs. WT | -0.4±0.2 | -1.3 to 0.5 | 0.21 | -0.5±0.04 | -0.7 to -0.3 | 0.008 |
| *acs* octuple vs. line #6-2 | -65.7±30.4 | -197.6 to 65.2 | 0.16 | -22.0±12.1 | -74.1 to 30.2 | 0.21 |
| *acs* octuple vs. line #7-1 | -52 ±23.8 | -154.5 to 50.6 | 0.01 | -17.3±8.3 | -53.3 to 18.6 | 0.17 |
| *acs* octuple vs. line #5-6 | -191.3±4.1 | -243.2 to -139.4 | 0.16 | -77.4±0.04 | -78 to -76.9 | 0.0003 |

**Table S6 qRT-PCR of *ACS8* and *ACS11* genes in the *GCD1::ACS8/11* transgenic lines.** The table shows ACS8 and ACS11 gene expression levels in the acs octuple mutant compared to WT and the C GCD1::ACS8/11. The tables present the relative ACS8 and ACS11 gene expression means difference, 95% confidence intervals, and P-value of the effect size (PES) of each separate comparison. The mean difference presents the averaged expression level in the acs octuple mutant compared to each line. The model parameters were analyzed as repeated measurements using two-way ANOVA followed by Fisher's LSD test.

| **Table S6** | **Mean difference ± SE** | **95% CI of diff.** | $P_{ES}$ | **Mean difference ± SE** | **95% CI of diff.** | $P_{ES}$ |
|---|---|---|---|---|---|---|
| *GCD1::ACS8/11* | ***ACS8*** | | | ***ACS11*** | | |
| *acs* octuple vs. WT | -0.53 ± 0.13 | -0.9 to -0.1 | 0.02 | -0.49 ± 0.18 | -1 to 0.1 | 0.07 |
| *acs* octuple vs. line #8-5 | -529.6 ± 78.15 | -865.9 to -193.3 | 0.02 | -2136 ± 589.9 | -4013 to -258.6 | 0.04 |
| *acs* octuple vs. line #10-1 | -16.72 ± 3.6 | -32.4 to -1.1 | 0.04 | -18.44 ± 3.16 | -32 to -4.8 | 0.03 |
| *acs* octuple vs. line #6-2 | -7.8 ± 3.04 | -17.5 to 1.9 | 0.08 | -3.531 ± 0.41 | -4.8 to -2.2 | 0.003 |

**Methods S1** ***Schematics of DNA constructs and expression promoters.*** ACS8/11 tissue-specific complementation constructs. (A) All constructs share a common foundational design for expressing a modified mutated version that complements the 2 genes of (a) ACS8 and (b) ACS11. (c) The green/yellow fluorescent protein - Neon green expression cassette. All constructs possess (d) cassette for screening, expressing the Venus yellow fluorescent protein under the CRU3 seed coat protein promoter. (B) All three genes are cloned in three different cassettes and use the same expression promoter in each construct: either 1) the guard cell-specific promoter; GC1 2) the spongy mesophyll-specific promoter, CORi3 3) the palisade mesophyll-specific promoter, IQD22 (PEG1), and 4) the whole-plant expression promoter, UBQ10.

**(A)**

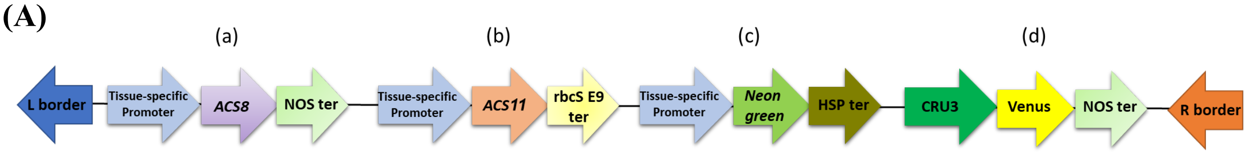


**(B)**

**<u>Expression Promoters</u>**

1) GC1 *(GCD1)* - a guard cell-specific expressing

2) CORi3 *(CORI3)* - a spongy mesophyll-specific

3) IQD22 *(PEG1)* - a palisade mesophyll-specif*ic*

4) UBIQUITIN 10 (UBQ10) - The whole-plant expression promoter
